# NPannotator: a genome- and chemistry- constrained automation for type I polyketide synthase pathway elucidation

**DOI:** 10.64898/2026.04.06.712324

**Authors:** Yash Chainani, Andre Cornman, Yunha Hwang

## Abstract

Natural products (NPs) are structurally diverse bioactive compounds whose biosynthesis is encoded within biosynthetic gene clusters (BGCs). Although databases such as the *Minimum Information about a Biosynthetic Gene Cluster* (MiBIG) repository now catalog thousands of experimentally validated NP structures, the full biosynthetic pathway connecting individual domain sequences to specific chemical features on final NP structures remains largely unannotated. This gap is especially pronounced for type I polyketide synthases (PKSs). These are modular assembly lines in which multiple enzymatic domains work in concert to condense acyl-CoA building blocks into complex polyketide scaffolds. Within these systems, acyltransferase (AT) domains govern which starter and extender units are incorporated at each elongation step, yet the substrate specificities of AT domains are known for only a fraction of cataloged clusters. Moreover, the catalytic order of genes encoding PKS modules is not immediately apparent from existing database entries, leaving the correct module ordering for observed product structures uncertain. Here, we present NPannotator, an automated, genomic context-aware cheminformatics pipeline that infers both the catalytic ordering of PKS domains and the substrate specificities of a given PKS’s AT domains. NPannotator loads a precomputed database of synthetically generated polyketide backbones, iteratively replaces default malonyl-CoA substrates with candidate starter and extender units via SMARTS-based substructure matching against the target NP, and selects the arrangement that maximizes chemical similarity. When benchmarked on the type I PKSs annotated within the expert-reviewed ClusterCAD dataset, NPannotator recovered 62.0% of both correct gene orderings and AT substrate annotations, and achieved 80.0% accuracy on gene ordering alone. By bridging gene-level architecture with chemical outcomes, NPannotator represents a step toward systematically decoding how protein sequence and genomic organization encode chemical structure in the world of natural products.

## 1. Introduction

Biosynthetic gene clusters (BGCs) are tightly co-regulated sets of genes found across microbial, fungal, and plant genomes^1–4^ that often encode large, multifunctional enzymes such as polyketide synthases (PKSs)^5–9^ and non-ribosomal peptide synthetases (NRPSs)^10–13^, which synthesize structurally complex natural products (NPs)^14^. These modular enzymes condense monomeric building blocks, acyl-CoA derivatives for PKSs^7–9^ and amino acids for NRPSs^11^, into therapeutically valuable compounds. Many NPs possess potent bioactive properties, and a substantial proportion of FDA-approved drugs are derived from NP structures or their chemical scaffolds^14–18^.

Yet this translates to a massive gap in our understanding: the antiSMASH database contains over 250,000 BGC entries, but only about 3,000 have experimentally verified structures in the Minimum Information about a Biosynthetic Gene Cluster (MiBIG) database^19,20^. Only 1% of the “chemical dark matter” has been uncovered. The disparity is staggering in the Joint Genome Institute’s Secondary Metabolism Collaboratory^21^ (SMC), which catalogs 1.2 million BGCs but comparatively few confirmed structures. Meanwhile, chemistry databases like the Natural Products Atlas^22^ and COCONUT^23^ contain thousands of natural product structures, yet their biosynthetic origins remain a mystery. Linking BGC sequences to final chemical structures remains one of the central challenges in natural product discovery^21,23–26^.

Previous efforts have approached the sequence-to-structure challenge from several complementary directions^19,26^. Rule-based approaches, such as antiSMASH^19^ and PRISM^27,28^, rely on curated biosynthetic heuristics and domain-specific motifs to generate rough predictions of possible chemical scaffolds, but these methods are limited by the completeness of known rules and cannot easily generalize to novel chemistry or sequences. For example, antiSMASH uses profile HMMs^19^ to identify core biosynthetic enzymes (e.g., PKS ketosynthases, NRPS adenylation and condensation domains) and co-occurrence rules to define cluster boundaries, and then applies simple assembly logic, such as the collinearity of modules in NRPSs or the predicted number of polyketide extension cycles based on the Chain Length Factor (CLF) phylogeny to propose core chains and scaffold backbones. It also encodes motif-based heuristics for tailoring enzyme classes (e.g., methyltransferase active site patterns or cyclase/ regioselectivity clades) to guess likely modifications and ring closures in type II polyketide scaffolds^12^. PRISM similarly leverages libraries of HMMs and manually curated biosynthetic reaction rules to assemble combinatorial chemical graph representations that reflect all biosynthetically plausible products consistent with a cluster’s enzyme repertoire. PRISM’s predictions can include specific features such as alternative starter units, predicted monomer sequences for NRPSs, and enumerated virtual tailoring reactions based on known enzyme specifies^27,28^.

These heuristics allow for the generation of rough core scaffold hypotheses (e.g., predicted NRPS sequences or polyketide backbones with putative tailoring patterns) but inherently rely on existing knowledge and cannot fully capture novel biosynthesis pathways. Domain-level predictors, including substrate classifiers for AT and A domains^29,30^, have used profile HMMs or sequence similarity to annotate monomer specificities, yet these often lack the resolution to capture subtle sequence determinants of substrate choice. Separately, methods such as PKSpop^31^ and DDAP^32^ have addressed the assembly line ordering problem by predicting docking domain interactions between PKS subunits, but these methods do not predict AT substrate specificity and thus address only one component of the sequence-to-structure mapping. While computational platforms for engineering chimeric PKSs^33^ have emerged from well-characterized pathways such as the methymycin/pikromycin system^34,35^, systematically discovering and annotating existing BGC-structure relationships is an equally important problem that remains unaddressed.

Here, we tackle the problem of mapping BGC sequences to resulting chemical structure by introducing NPannotator, a computational framework designed to bridge the gap between BGC sequences and their corresponding NP structures by elucidating the catalytic order of modules and predicting the substrate specificity of AT domains in type I PKSs. Given a NP molecular structure and its corresponding BGC sequence entry (e.g. from MiBIG^20^), NPannotator extracts and applies a set of genomic constraints to reconstruct plausible domain/ module organizations. It then evaluates these candidate reconstructions by comparing the predicted domain-substrate assignments against known biosynthetic logic, selecting the configuration that best matches the target structure. NPannotator explicitly tests for concordance between predicted and experimentally validated domain orders and substrate profiles, thereby enabling direct assessment of annotation accuracy. We validate NPannotator against the ClusterCAD dataset^36^, which provides expert-reviewed domain orderings and substrate annotations for 183 PKS, NRPS, and hybrid clusters, offering a rigorous benchmark for evaluating annotation tools. By integrating genomic organization, biosynthetic constraints, and structural matching in a single pipeline, NPannotator provides a scalable and generalizable approach for linking NP structures to their biosynthetic gene clusters.

**Figure 1.**
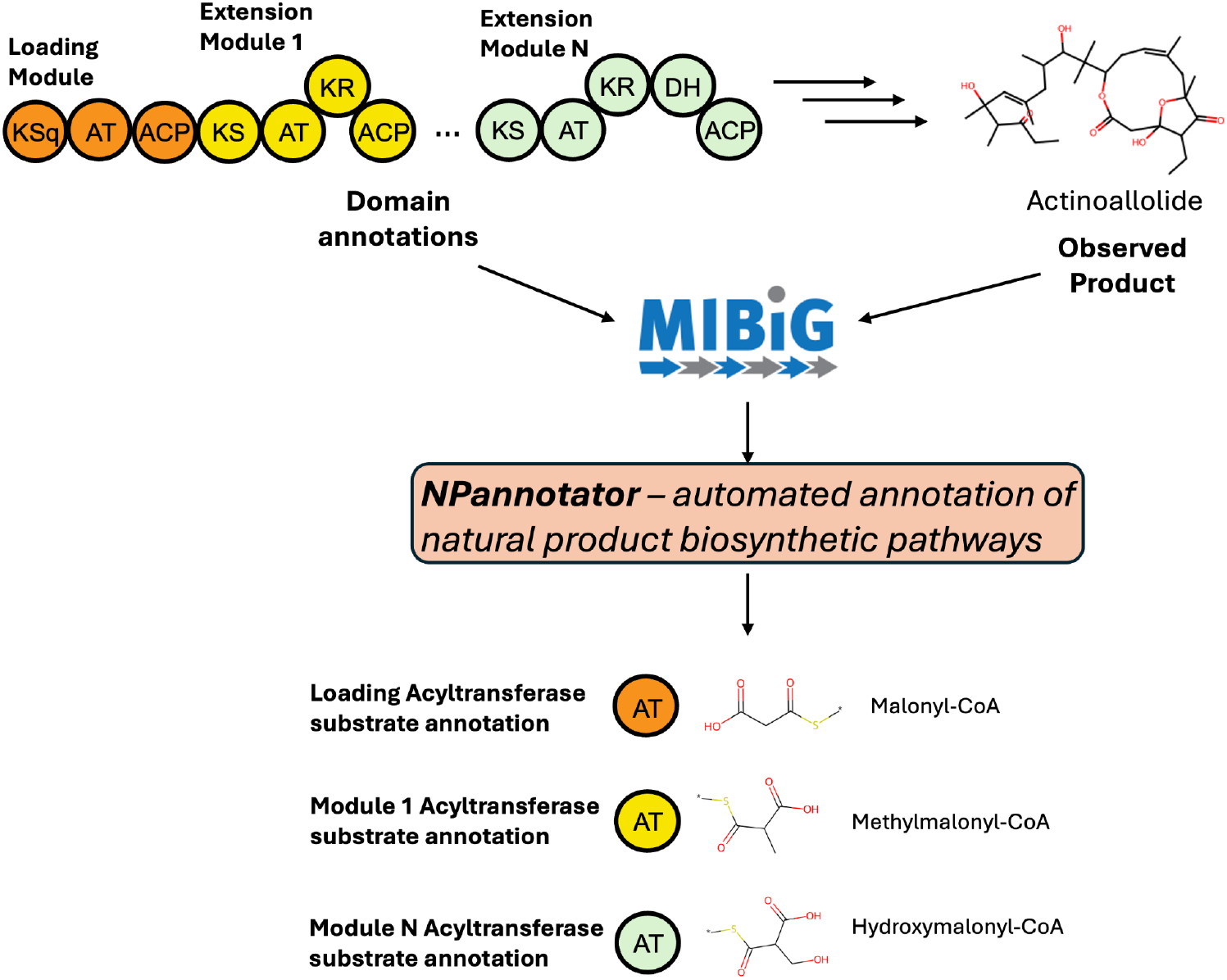
NPannotator elucidates the intermediate substrates involved in biosynthesis of experimentally validated type I polyketide synthase gene clusters. NPannotator provides automated annotation of natural product biosynthetic pathways. Biosynthetic gene clusters deposited within the Minimum Information for a Biosynthetic Gene Cluster (MIBiG) database only comprise domain annotations and final product annotations but the intermediate substrate annotations required to assemble the final natural product are unknown. NPannotator provides these annotations by iteratively replacing the default malonyl-CoA substrate with different intermediate substrates until a scaffold that is most chemically similar to the target natural product is reached.

## 2. Methods

### 2.1 Modules in NPannotator

NPannotator is a fully open-source package that is written completely in the Python programming language. It comprises two key modules: (1) the MiBIG parser and (2) the annotator. The input to the MiBIG parser module is a .json file representing a type I PKS BGC and the output is a list of enzymatic domains and their genomic organization, i.e., which domains are co-located on the same gene. The enzymatic domains that the parser module extracts will belong to either one of the following types: “PKS_AT” (acyltransferase domain), “PKS_KS” (ketosynthase domain), “PKS_KR” (ketoreductase domain), “PKS_DH” / “PKS_DHt” (dehydratase domains, the latter being a variant), “PKS_ER” (enoylreductase domain), “Thioesterase” / “TD” (termination domain), “PKS_Docking_Nterm”, or “PKS_Docking_Cterm” (inter-module or inter-domain linkers). The MiBIG parser also extracts protein sequences for each domain. These extracted domains will then be used to initialize corresponding reaction rules from the biosynthetic cluster simulator (BCS) module of the previously released RetroTide package for designing type I PKS chimeras^33^. For instance, the “PKS_AT” domain extracted from a given MiBIG file will map to a bcs.AT reaction template while a “PKS_KR” domain will map to a bcs.KR template. Reaction rules in the bcs package are chemical templates written in the SMiles Arbitrary Target Specification (SMARTS) chemical language first developed by Daylight Chemical Information Systems, Inc. These reaction rules represent the enzymatic transformation catalyzed by each type I PKS domain. If a “CAL” domain is present representing a coenzyme A (CoA) ligase domain, then this is initially read in as an acyltransferase domain since both domains are functionally similar in terms of catalyzing the acylation of an acyl-CoA building block (e.g., malonyl-CoA or methylmalonyl-CoA) onto the loading module’s acyl carrier protein (ACP) domain. We note that the “PKS_ACP” domain representing the ACP domain in MiBIG is not read in because its function of ensuring the growing polyketide chain remains tethered to the full PKS assembly is already implicitly captured within RetroTide’s bcs module. Further, the “PKS_KS” domain does not have a corresponding reaction template because the Claisen condensation reaction catalyzed by this domain is also implicitly captured within RetroTide when two module objects, each representing a list of domain objects, are concatenated together.

The outputs from the MiBIG parser module, which comprise (1) a list of enzymatic domains, (2) their corresponding genomic arrangement, and (3) the chemical structure of the final compound, will then be fed as part of the inputs into the annotator module. The chemical structure of the final compound will be captured in terms of its Simplified Molecular Input Linear Entry System (SMILES) string. The annotator module will later convert this SMILES string into a mol object using the chemformatics package RDKit in order to sanitize this string and remove any associated stereochemistry information.

### 2.2 Identification of loading module, terminal module, and corresponding genes

In addition to extracting the genomic constraints that describe which enzymatic domains are co-located on the same gene, the MiBIG parser module within our NPannotator package also determines the boundaries of a given PKS assembly by identifying the genes encoding for the initial loading module and final termination module. In type I PKSs, the loading module - characterized by the presence of a loading ACP domain - appears at the start of a gene cluster and contains only a C-terminal docking domain (“PKS_Docking_Cterm”) without an N-terminal docking domain (“PKS_Docking_Nterm”). All modules encoded on this gene are therefore positioned in the left-to-right catalytic order. Conversely, the final catalytic module in a type I PKS is immediately followed by a termination domain (“Thioesterase” or “TD”), allowing both the last module and its corresponding gene to be unambiguously identified. For example, given the BGC of Pikromycin^34,35^ (MiBIG accession ID: BGC0000094.1) with the genes pikAI, pikAII, pikAIII, and pikAIV, the MiBIG parser module in NPannotator can identify that the gene pikAI and its associated modules (1 loading module and 2 extension modules) must appear first given the lack of a N-terminal docking domain. Similarly, the gene pikAIV and its associated modules must appear last given the presence of a thioesterase (TE) domain.

### 2.3 Enumeration of possible genomic configurations by the annotator module

With a list of genes, their constituent modules, and the catalytic domains within each module extracted by the MiBIG parser module of NPannotator, the annotator module now proceeds to enumerate all possible arrangements of the biosynthetic assembly line that comply with the extracted genomic constraints. Since the start and end genes have been identified thus far, these are always held fixed in the first and last positions, respectively while the remaining internal genes - which encode for modules positioned between the loading and terminal modules - are permuted around in all possible orders.

For each permutation, NPannotator concatenates the modules from the start gene, the permitted internal genes, and the end gene to produce a hypothetical biosynthetic sequence of enzymatic domains/ modules. This hypothetical sequence preserves the left-to-right order of modules within internal genes as recorded in each MiBIG entry, thereby ensuring that intragenic domain arrangements remain intact while allowing for flexibility in the intergenic ordering of modules. The enumeration step therefore explores all plausible gene-level organizations that could correspond to the true catalytic order in the native biosynthetic pathway.

### 2.4 Precomputing a database of candidate chemical scaffolds

In order to accelerate downstream annotation, NPannotator uses a precomputed database of synthetically generated type I PKS chemical scaffolds. These scaffolds were generated by running the bcs module of the RetroTide package forwards for 11 extension modules using only malonyl-CoA as both the initial loading and subsequent extension units^33^. Since malonyl-CoA is the simplest repeat unit with no side chain atoms and only two carbon atoms, it serves as a minimal baseline backbone that preserves only the carbon–oxygen framework and off-backbone functional groups dictated by the KR, DH, and ER domains.

Each scaffold in this database corresponds to a unique combination of module architectures up to 11 extension modules in length. The bcs module applies the corresponding SMARTS-based reaction templates for each domain in sequence, starting from a malonyl-CoA loading unit to construct the linear polyketide chain *in silico*. For every generated scaffold, the resulting chemical structure is stored as a SMILES string along with metadata describing the ordered list of domains and the total number of AT domains.

When attempting to annotate a target BGC, NPannotator first filters the precomputed database to retain only those scaffolds that contain the same number of AT domains as the target PKS. Since the AT count is independent of module order, this step immediately removes length-incompatible designs from consideration. The filtered set is then pared down further to retain only those scaffolds that share the same set of module architectures as the target (excluding the loading module), where each module is represented as an unordered set of domains. This ensures that the candidate scaffolds match both the total chain length and the per-module domain composition of the target PKS while leaving the catalytic order open for evaluation.

### 2.5 Extraction of target molecule’s carbon and oxygen scaffold

Here we derive a carbon–oxygen (CO) scaffold from the target RDKit molecule by isolating the largest contiguous carbon framework and appending only directly bonded oxygens. Concretely, we collect all carbon atoms and carbon–carbon bonds to build a carbon-only graph, retain its largest connected component, and substructure-match that component back to the original molecule to recover the corresponding atom indices. We then augment this backbone with any immediate oxygen neighbors of those carbons, reconstruct a new molecule containing only these C and O atoms and their interconnecting bonds, and sanitize the result; if sanitization fails, we conservatively return the original molecule with a warning. The resulting CO scaffold preserves backbone connectivity and proximal carbonyl/ether features while discarding distal substituents and stereochemistry, yielding a fast, robust template for the starter down-selection step. The CO scaffold is also used to detect macrolactone rings via a SMARTS pattern matching for cyclic esters, which determines whether the offloading step applies cyclization or thiolysis.

### 2.6 Optimization of starter acyl-CoA unit

NPannotator narrows the search space by selecting plausible starter acyl-CoA units for the loading module. Our library includes approximately 25 different starter unit types, encompassing common starters (acetyl-CoA, propionyl-CoA, benzoyl-CoA) and specialized ones (AHBA in both oxidized and reduced forms, DHCH, CHC-CoA, and cyclopentene-based starters). When a KSq (decarboxylating ketosynthase) domain is present, only non-monocarboxylic starters are evaluated. Candidate starters are down-selected using SMARTS-based substructure templates matched against both the full target molecule and its carbon–oxygen (CO) scaffold, and the smaller set is retained. For each retained starter acyl-CoA unit, NPannotator regenerates the cluster with the new loading unit, simulates product formation using RetroTide, and applies one of four offloading mechanisms: thiolysis (releasing free carboxylic acid), macrocyclization (forming a lactone), full reduction, or reduction to aldehyde. When a macrolactone is detected, only products matching the target’s largest ring size pass through. Candidate products are filtered based on genomic constraints and on their agreement with the functional-group composition of the target CO scaffold.

### 2.7 Optimization of extender acyl-CoA units

Following starter optimization, NPannotator selects extender acyl-CoA units for each AT domain. Five extender types are tested: malonyl-CoA, methylmalonyl-CoA, ethylmalonyl-CoA, methoxymalonyl-CoA, and allylmalonyl-CoA. SMARTS patterns are used to detect and count each extender type in the target molecule. Special handling for methylmalonyl-CoA includes a lactone-adjacent methyl group detection pattern that adjusts the expected methylmalonyl count. The scoring uses a two-phase approach: first filtering by off-backbone element ratios, then ranking by stereo-agnostic maximum common substructure (MCS)-based Tanimoto similarity. All valid extender combinations are enumerated across extension modules, clusters regenerated, and products simulated in parallel using Python’s multiprocessing.Pool for computational efficiency. Candidate products are then filtered by genomic constraints, agreement with the target scaffold, and extender usage consistency, and are finally ranked by chemical similarity to the target. The highest-ranked configuration is reported as the predicted gene ordering and AT assignments.

## 3. Results

### 3.1 Architecture of NPannotator

NPannotator begins with a precomputed database of malonyl-CoA–initiated scaffolds and uses substructure templates to identify plausible starting substrates for the target biosynthetic gene cluster (BGC). Both the full target natural product and its derived carbon–oxygen (CO) scaffold (see Methods 2.5) are queried, and the smaller of the two resulting sets (typically 5–15 candidates from an initial library of ∼30 starters) is retained. For each retained starter, NPannotator rebuilds the cluster, substituting the new loading unit into the first AT domain, and simulates the resulting product using RetroTide’s bcs forward synthesis module^33^. Products are offloaded according to the target’s macrocyclization state: if the target CO scaffold contains a lactone, NPannotator attempts cyclization to the correct ring size; otherwise, it applies a reductive release via a thiolysis reaction to form a carboxylic acid.

To filter these products, NPannotator compares their offloaded backbones against the target CO scaffold using maximum common substructure (MCS) analysis. Functional groups such as carbonyls, hydroxyls, and alkenes are enumerated in the MCS, and a non-C-C single-bond ratio is calculated as the sum of these group counts normalized by the target. Only candidates whose ratios are closest to 1.0 (indicating faithful reproduction of the target scaffold’s functional-group density) are retained, and candidates that violate genomic ordering constraints are discarded.

From the remaining starter-optimized clusters, NPannotator next determines which extender units (e.g., malonyl-CoA, methylmalonyl-CoA, ethylmalonyl-CoA) are most consistent with the target. This is achieved by template matching against both the target molecule and its CO scaffold, yielding a bounded set of extenders (usually 2 to 4 per case). For each extender type, upper limits on allowable counts are inferred, with special handling of methylmalonyl-CoA: NPannotator enforces soft lower and upper bounds around the predicted count to avoid under- or overestimating branching frequency.

All valid combinations of extenders are then enumerated across the extension modules of each candidate cluster. For example, a cluster with six AT domains and three extender types may yield hundreds of possible assignments, all of which are generated and simulated in parallel. Each new product is offloaded as before, and genomic constraints are re-enforced.

To prioritize among the resulting candidates, NPannotator evaluates each offloaded product against the target CO scaffold along three axes:

1. Functional-group preservation: using maximum common substructure (MCS)-derived ratios of carbonyls, hydroxyls, and alkenes to ensure off-backbone chemistry matches the target.
2. Extender unit consistency: comparing the realized counts of malonyl-, methylmalonyl-, and ethylmalonyl-CoA units to the target-inferred maxima. Products are sorted first by methylmalonyl ratio, then malonyl ratio, then ethylmalonyl ratio.
3. Structural similarity: computing a stereochemistry-agnostic MCS-based Tanimoto similarity score, as implemented in RetroTide^33^, between each candidate scaffold and the target CO scaffold.

Only candidates achieving the highest methylmalonyl ratio are retained, and among these, those with the highest ethylmalonyl ratio are further prioritized. The final ranking is determined by similarity score, and the top-scoring cluster is selected as NPannotator’s prediction of the most likely gene ordering and AT substrate assignment for the target BGC.

In practice, this staged procedure rapidly reduces the combinatorial search space. For instance, from an initial library of tens of thousands of precomputed malonyl-CoA backbones, NPannotator typically narrows the candidate set to fewer than 20 starter variants and a few hundred extender combinations, which can be evaluated within minutes on a multicore CPU. This systematic pruning, guided by structural matching at every step, allows NPannotator to produce accurate annotations even in the face of incomplete or ambiguous genomic information.

### 3.2 Worked example for annotating Pikromycin

The following walkthrough demonstrates how NPannotator processes the pikromycin biosynthetic gene cluster (MiBIG accession BGC0000094.1). The MiBIG parser reads the cluster annotation and extracts four genes: *pikAI, pikAII, pikAIII*, and *pikAIV*. The parser identifies *pikAI* as the start gene (since it possesses only a C-terminal docking domain with no N-terminal docking domain) containing one loading module and two extension modules. The gene pikAIV is identified as the terminal gene because it harbors a thioesterase domain, the enzyme responsible for catalyzing the release of the fully elongated polyketide chain from the PKS assembly line. Genomic constraints are extracted: modules on the same gene must remain contiguous in the catalytic order. The annotator loads the precomputed scaffolds database and filters for clusters with seven AT domains (matching pikromycin) and matching domain architectures. Starter down-selection via SMARTS templates identifies malonyl-CoA as compatible based on substructure matching against pikromycin’s target structure. Clusters are regenerated with this starter, products computed and offloaded via macrocyclization (pikromycin contains a macrolactone), and candidates filtered by ring size and off-backbone element ratios. Extender iteration detects methylmalonyl-CoA and malonyl-CoA as relevant extenders, and all valid combinations are enumerated and evaluated in parallel. Final ranking by MCS-based Tanimoto similarity selects the top configuration, recovering the correct gene order (pikAI → pikAIII → pikAII → pikAIV) and substrate assignments.

### 3.3 Recovery rate of gene cluster annotations provided by a high quality dataset

We benchmarked NPannotator against the ClusterCAD dataset, which comprises 183 expert-reviewed PKSs, NRPSs, and PKS–NRPS hybrids with experimentally validated domain orders and substrate annotations. This dataset provides a uniquely high-quality reference for evaluating annotation tools, since each entry encodes not only a validated natural product structure but also domain-level and substrate-level assignments curated by experts. Because NPannotator currently targets type I PKSs and supports assembly lines of up to 11 extension modules (n = 50 clusters), we restricted our evaluation to the subset of ClusterCAD entries meeting these criteria. We tested NPannotator in two modes: (i) requiring the recovery of both gene ordering and AT substrate assignments, and (ii) requiring only gene ordering without substrate-level specificity. In the more stringent task (Domains + Substrates), NPannotator correctly recovered annotations for 62.0% of clusters. When only domain ordering was considered, accuracy increased to 80.0% of clusters. By comparison, a random baseline in which substrates as well as module orderings are randomly chosen achieves a mean accuracy of just 8.0% for domain ordering alone and 0.0% for both substrate assignment and the joint task, thereby confirming that NPannotator’s performance is far above random chance (Figure 2).

**Figure 2.**
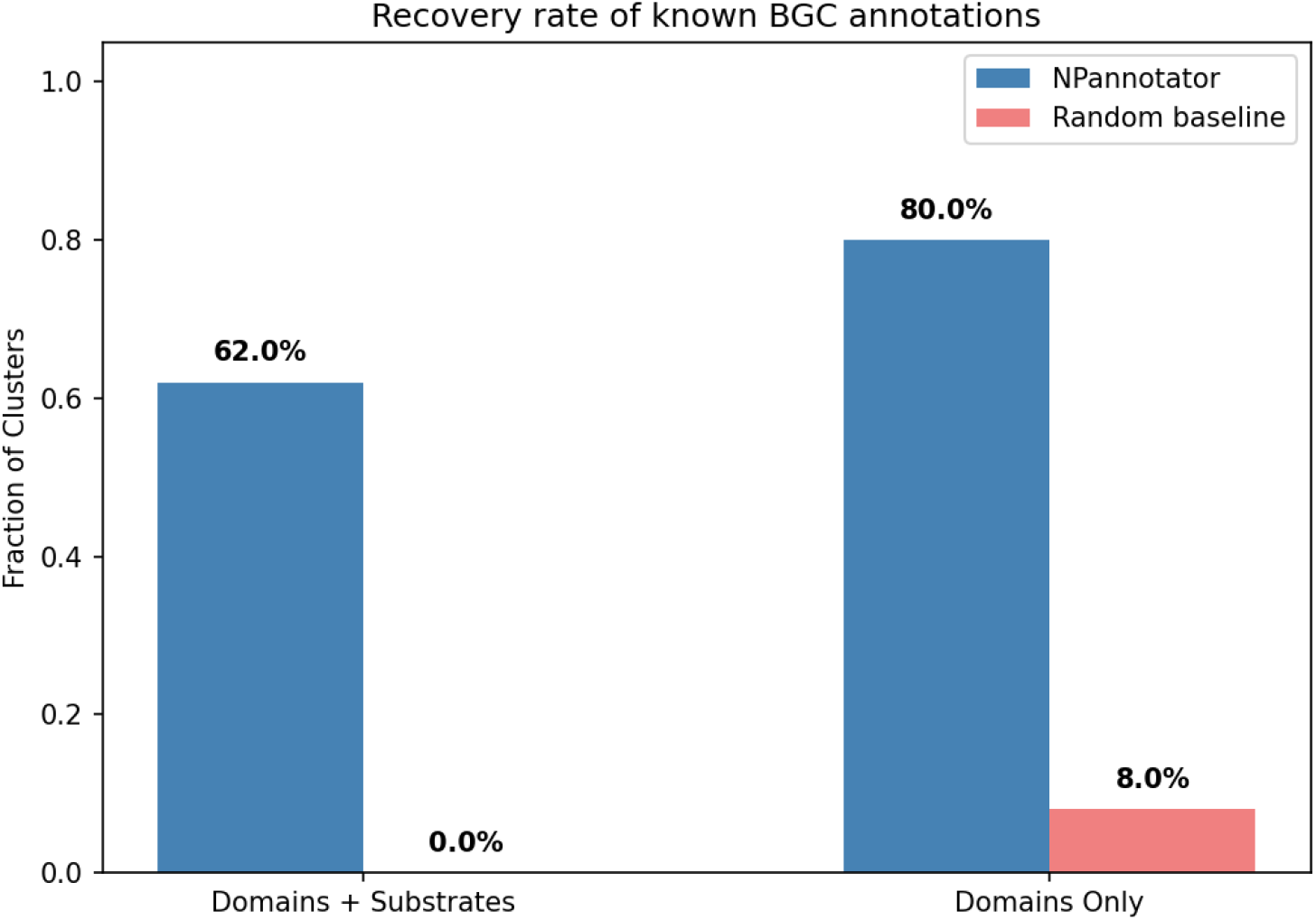
Recovery rate of NPannotator annotations on the ClusterCAD dataset. Blue bars show the fraction of clusters correctly annotated by NPannotator when requiring both domain ordering and AT substrate assignments (“Domains + Substrates,” 62.0 %) and when only domain ordering is considered (“Domains Only,” 80.0 %). Red bars show a random baseline in which domain orderings and substrates were selected at random. For domain ordering, the loading module is fixed and only the N−1 extension modules are permuted, accounting for identical domain architectures; the mean probability of a correct random ordering is 8.0%. For substrate assignment, the probability of randomly guessing all substrates correctly is 0.0%. The joint Domains + Substrates baseline is similarly 0.0%. A random choice baseline is used as the comparison because no existing tool jointly predicts both catalytic gene order and AT substrate specificity: assembly line ordering methods such as DDAP^32^ and PKSpop^31^ infer gene order via docking domain interactions but do not predict substrate assignments, while tools that address substrate specificity (e.g., antiSMASH’s pHMM-based AT predictions) assume gene order is already known, making none directly comparable to NPannotator’s end-to-end prediction task.

These results highlight two key points. First, NPannotator is able to reconstruct the catalytic sequence of modules with high fidelity, even when gene order is not explicitly encoded in MiBIG. Second, although extending this accuracy to AT substrate assignments introduces additional challenges, owing to the subtlety of sequence-to-substrate determinants and the diversity of acyl-CoA building blocks, NPannotator still performs robustly, recovering the majority of correct assignments. Importantly, both performance levels significantly exceed what would be expected by random assignment, demonstrating that the pipeline’s combined use of genomic constraints, precomputed scaffolds, and chemical similarity scoring provides meaningful predictive power.

### 3.4 Limitations of NPannotator

A key limitation of NPannotator is its reliance on chemical similarity to a known product structure, which can be confounded by post-PKS tailoring modifications. After the PKS assembly line releases a polyketide chain, tailoring enzymes frequently remodel the scaffold in ways that obscure its biosynthetic origin: cytochrome P450 monooxygenases introduce hydroxyl groups at specific backbone positions, glycosyltransferases attach sugar moieties (as seen in erythromycin and pikromycin), methyltransferases add methyl groups that can mimic methylmalonyl-CoA-derived branching, and oxidative enzymes can rearrange ring systems entirely. NPannotator partially mitigates the effect of P450 hydroxylations by using a SMARTS-based heuristic to identify and subtract likely post-PKS hydroxyl additions from its off-backbone element counts, but this correction is imperfect and cannot reliably distinguish PKS-intrinsic hydroxyls from those introduced by tailoring. More broadly, because NPannotator ranks candidate scaffolds by their chemical similarity to the target structure, heavily tailored products can mislead the scoring: backbone-consistent candidates may be ranked below incorrect alternatives that coincidentally share more substructural features with the final, decorated molecule. NPannotator addresses this in part by comparing against the carbon–oxygen (CO) scaffold rather than the full molecule, but even the CO scaffold can diverge substantially from the true PKS product when tailoring is extensive.

Figure 3 illustrates this challenge concretely. When NPannotator ranks candidate scaffolds for a macrocyclic polyketide target, the true PKS-derived backbone is placed fifth, behind four alternatives that achieve higher atom-pair similarity, Tanimoto similarity, and MCS overlap scores. Each of these higher-ranked candidates shares more substructural features with the final, tailored natural product, but not with the actual biosynthetic intermediate from which it was derived. The mis-ranking arises precisely because post-PKS modifications shift the molecule’s similarity profile away from the underlying polyketide skeleton and toward decorated structures that were never part of the PKS assembly line.

**Figure 3.**
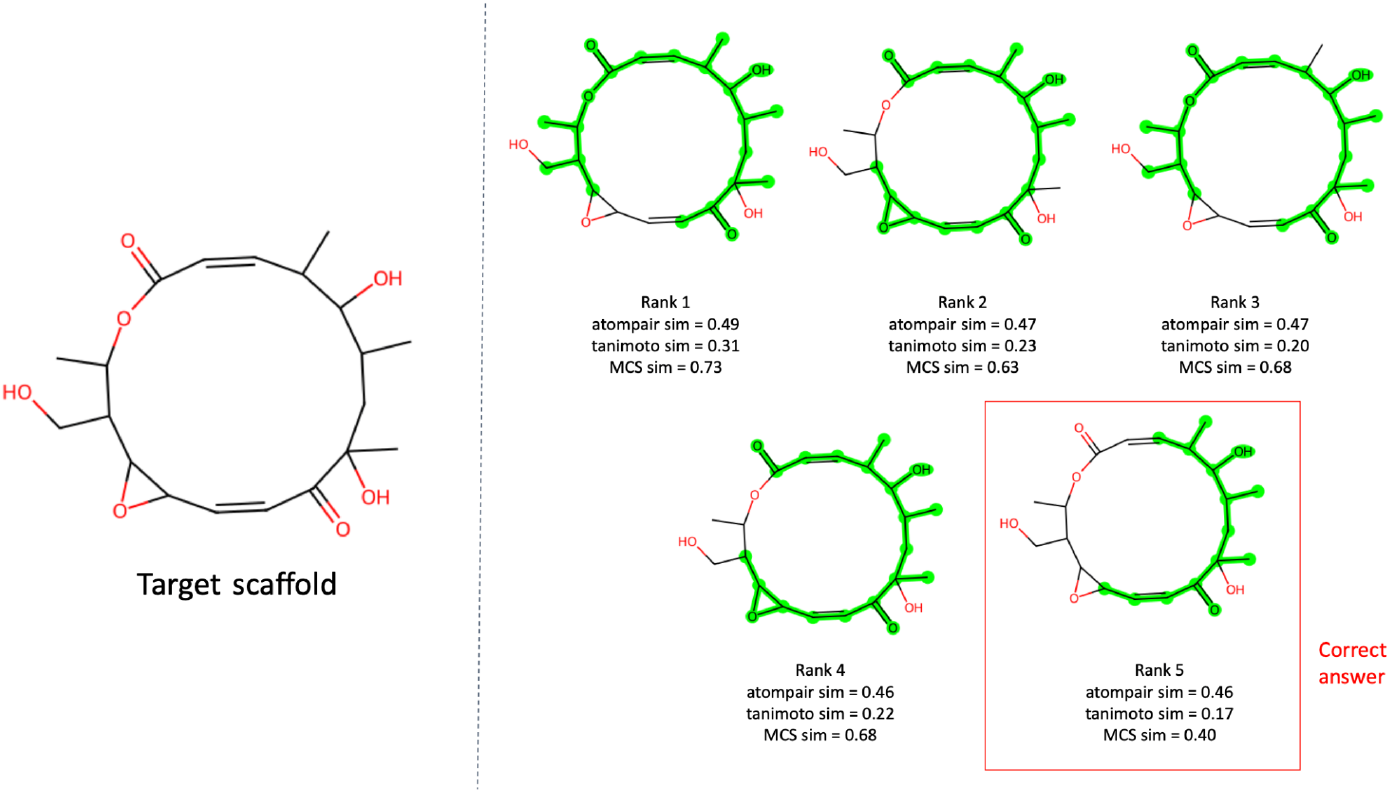
Standard chemical similarity metrics can misidentify true polyketide backbones due to post-PKS tailoring modifications. A macrocyclic polyketide natural product is shown alongside candidate biosynthetic scaffolds ranked by standard chemical similarity metrics, including atom-pair similarity, Tanimoto similarity, and maximum common substructure (MCS). Although NPannotator ranks candidates based on these metrics, the true PKS-derived backbone (highlighted) is ranked fifth while higher-ranked candidates (ranks 1-4) achieve greater similarity scores despite deviating from the correct biosynthetic intermediate. This misidentification arises because post-PKS tailoring reactions, such as oxidations, hydroxylations, or ring arrangements, can substantially alter the final natural product structure, shifting its similarity profile away from the underlying PKS-encoded scaffold. This example illustrates the limitations of relying on chemical similarity alone and motivates the incorporation of biosynthetic logic and tailoring-aware constraints to improve scaffold annotation.

This example underscores that chemical similarity, while effective for narrowing the combinatorial search space, is insufficient as a sole ranking criterion when tailoring is extensive. Addressing this limitation will require moving beyond static similarity comparisons toward approaches that account for the biosynthetic context of structural features. Promising directions include predicting which tailoring domains are present in a given BGC and penalizing candidate scaffolds that would require biosynthetically implausible post-PKS transformations to reach the observed product. More broadly, incorporating cheminformatic rules for common tailoring reactions, such as hydroxylations, glycosylations, methylations, and oxidative rearrangements, could allow NPannotator to reason backward from a decorated product to its most likely polyketide precursor, rather than comparing forward from candidate scaffolds to the final structure.

## 4. Conclusions and future work

In this work, we introduced NPannotator, a computational framework that bridges the gap between biosynthetic gene clusters and their encoded natural product structures. By systematically enumerating genomic constraints, simulating candidate polyketide backbones, and iteratively optimizing starter and extender unit assignments, NPannotator provides accurate predictions of both catalytic gene order and AT domain substrate specificity. Benchmarking against the high-quality ClusterCAD dataset^36^ demonstrated that NPannotator can recover 80.0% of gene orderings and 62.0% of full gene orderings with substrates—vastly exceeding an analytically computed random baseline of 8.0% and 0.0%, respectively—establishing its utility as a scalable and completely automated annotation tool for modular polyketide synthases.

Type I modular polyketide synthases were selected as an initial proof-of-concept system because they offer a well-defined, collinear relationship between genomic organization and chemical assembly logic, making them particularly amenable to cheminformatics-based pattern matching with various substrates. Their modular architecture, recurrent domain composition, and existing biochemical characterization provide a strong foundation for testing sequence-to-chemical mapping approaches.

At the same time, our analysis revealed key challenges that remain for future development. Chief among these is the limitation of chemical similarity metrics, which can mis-rank correct backbones in cases where post-PKS tailoring enzymes introduce significant structural modifications. Addressing this limitation represents an important direction for future development. Promising avenues include: (1) incorporating cheminformatics rules for common post-PKS enzyme decorations, such as hydroxylations, glycosylations, and methylations; (2) integrating machine learning models trained on paired BGC-structure data to learn sequence-to-substrate and sequence-to-modification relationships directly; and (3) introducing reaction-context penalties that downweight candidates requiring unlikely post-PKS tailoring transformations.

Future extensions of NPannotator will focus on broadening its biosynthetic scope and improving scalability. In parallel, the cheminformatics-based annotations generated by NPannotator can serve as high-quality training labels for machine learning models that predict AT substrate specificity directly from protein sequence. Recent advances in protein language models, including the evolutionary scale family of language models^37^ (ESM) and genomic language models^38,39^ (gLM2) produce rich sequence embeddings that capture evolutionary and functional signals in AT domain sequences. By pairing NPannotator-derived substrate labels with these embeddings, it becomes possible to train classifiers (such as logistic regression or k-nearest neighbors) that generalize substrate predictions to uncharacterized AT sequences without requiring a known product structure. This complementary sequence-based approach would extend NPannotator’s reach from the approximately 3,000 BGCs with known products to the hundreds of thousands of unannotated gene clusters in public databases. These include expanding support to additional PKS subclasses (e.g., iterative and trans-AT PKSs), hybrid PKS-NRPS systems, and tailoring enzyme modules beyond core AT/KR/DH/ER chemistry. These advances will move NPannotator toward a more general and extensible sequence-to-chemistry inference framework, enabling systematic exploration of natural product biosynthetic diversity at scale.

Together, these extensions will enable NPannotator to scale beyond gold-standard datasets like ClusterCAD^36^ to more heterogeneous resources such as MiBIG and the JGI Secondary Metabolite Collaboratory^21^. More broadly, NPannotator represents one step toward a generalizable genome-to-chemistry mapping framework, helping to bridge the longstanding gap between biosynthetic gene cluster sequence data and the chemical structures they encode.

## Acknowledgements

This work is funded by the Gordon and Betty Moore Foundation through Grant GBMF13344 to Tatta Bio.

## Data and Code Availability

The NPannotator source code and associated data are publicly available at https://github.com/TattaBio/NPannotator.

